# MiRNAs differentially expressed in vegetative and reproductive organs of *Marchantia polymorpha* – insights into their expression pattern, gene structures and function

**DOI:** 10.1101/2023.11.30.569353

**Authors:** B Aggarwal, MW Karlowski, P Nuc, A Jarmolowski, Z Szweykowska-Kulinska, H Pietrykowska

## Abstract

MicroRNAs (miRNAs) regulate gene expression affecting a variety of plant developmental processes. The evolutionary position of *Marchantia polymorpha* makes it a significant model to understand miRNA-mediated gene regulatory pathways in plants. Previous studies focused on conserved miRNA-target mRNA modules showed their critical role in Marchantia development. Here, we demonstrate that differential expression of conserved miRNAs and their targets in selected organs of Marchantia additionally underlines their role in regulating fundamental developmental processes. The main aim of this study was to characterize selected liverwort-specific miRNAs, as there is a limited knowledge on their biogenesis, accumulation, targets, and function in Marchantia. We demonstrate their differential accumulation in vegetative and generative organs. We reveal that all liverwort-specific miRNAs examined are encoded by independent transcriptional units. MpmiR11737a, MpmiR11887 and MpmiR11796, annotated as being encoded within protein-encoding genes, have their own independent transcription start sites. The analysis of selected liverwort-specific miRNAs and their pri-miRNAs often reveal correlation in their levels, suggesting transcriptional regulation. However, MpmiR11796 shows a reverse correlation to its pri-miRNA level, suggesting post-transcriptional regulation. Moreover, we identify novel targets for selected liverwort-specific miRNAs and demonstrate an inverse correlation between their expression and miRNA accumulation. In the case of one miRNA precursor, we provide evidence that it encodes two functional miRNAs with two independent targets. Overall, our research sheds light on liverwort-specific miRNA gene structure, provides new data on their biogenesis and expression regulation. Furthermore, identifying their targets, we hypothesize the potential role of these miRNAs in early land plant development and functioning.

## Introduction

MicroRNAs (miRNAs) represent a class of endogenous, short (18-24nt in length), noncoding small RNA (sRNA) molecules whose expression is under tight control as they further direct the cleavage or cause translational repression of mRNAs encoding proteins responsible for plant development and responses to environmental cues (Jones-Rhoades et al., 2006; Chen, 2009; Luo et al., 2013; Spanudakis & Jackson, 2014; Hong & Jackson, 2015; Liu et al., 2018). Plant miRNAs are processed from *MIRNA* transcripts generated by RNA Polymerase II (Pol II). The primary transcript (pri-miRNA) bears the characteristics of typical Pol II transcript i.e., 5’-cap and 3’-poly(A) tail (Xie et al., 2005; Stepien et al., 2017; Wang et al., 2019). In the nucleus, pri-miRNAs undergo an initial cleavage performed by a RNase III enzyme Dicer-like 1 (DCL1), resulting in the release of shorter, stem-loop structured pre-miRNAs. In the second step, DCL1 again catalyzes endonucleolytic cleavage to further process the pre-miRNAs into miRNA/miRNA* duplexes. Subsequently, the miRNA/miRNA* duplexes are methylated at their 3’ ends by Hua enhancer 1 (HEN1) methylase (Chen, 2005; Bologna et al., 2009; Cuperus et al., 2010; Rogers & Chen, 2013; Dolata et al., 2018). These are then incorporated into ARGONAUTE 1 (AGO1) and exported into the cytoplasm where they perform their functions (Bologna et al., 2018; Zhu et al., 2022). Plant miRNAs are encoded by independent transcriptional units or are found within exons or introns of protein- or long noncoding RNA (lncRNA) genes. Splicing and alternative splicing of introns creates additional layer of post-transcriptional regulation of mature miRNA level. Also, alternative transcriptional start sites (TSS), alternative polyadenylation sites, pri-miRNA modifications (e.g., m6A), adoption of alternative stem-loop structure, selection of loop-to-base or base-to-loop processing mode of pri-miRNA or genes overlap may affect miRNA accumulation (Bielewicz et al., 2013; Li & Yu, 2021; Gonzalo et al., 2022; L. Zhang et al., 2022a; Bajczyk et al., 2023; Xu & Chen, 2023).

As compared to angiosperms, in which *MIR* genes are mostly found in the intergenic regions, most bioinformatically identified pre-miRNAs in *M. polymorpha* overlap with protein-coding genes. Nevertheless, independent transcriptional units encoding miRNAs were also found (Bowman et al., 2017; Pietrykowska et al., 2022). However, experimental validation of Marchantia *MIR* gene structures is missing. In this paper we show that selected Marchantia *MIR* genes that bioinformatically were found to overlap with protein-coding genes represent independent transcriptional units.

In angiosperms, at least eleven conserved miRNA families: miR156, miR159, miR160, miR164, miR166, miR167, miR169, miR172, miR319, miR390 and miR399, have been shown to affect plant development, including flowering time, flower development and seed production (Fahlgren et al., 2006; Jones-Rhoades et al., 2006; Palatnik et al., 2007; Yamaguchi et al., 2009; Liu & Vance, 2010; Nag & Jack, 2010; Hong & Jackson, 2015; Samad et al., 2017). In *M. polymorpha*, nine conserved miRNA families are present, which are found in all other land plants so far studied: miR160, miR166, miR171, miR319, miR390, miR529, miR408 and miR530/1030, miR536 (Lin et al., 2016; Tsuzuki et al., 2016; Bowman et al., 2017). To date, comparison of miRNA set in two liverwort species, *M. polymorpha* and *Pellia endiviifolia*, has revealed the presence of a very limited number of liverwort-specific miRNAs. These include three miRNA families: Pen-miR8163/MpmiR11737a/b, Pen-miR8185/MpmiR11889 and Pen-miR8170/MpmiR11865* (Tsuzuki et al., 2016; Bowman et al., 2017; Pietrykowska et al., 2022).

Thus far, the regulatory functions of several conserved, as well as liverwort-specific miRNAs, and their targets have been investigated in Marchantia. These studies have explored their roles in thallus development, phase transition, and sexual/asexual reproduction (Lin et al., 2016; Proust et al., 2016; Tsuzuki et al., 2016; Flores-Sandoval et al., 2018b; Honkanen et al., 2018; Tsuzuki et al., 2019; Thamm et al., 2020; Streubel et al., 2023). However, to gain a comprehensive understanding of the involvement of miRNAs and their targets in the intricate regulatory pathways that control Marchantia’s life cycle, comparative studies focused on these functions in liverworts and other basal plants would be essential.

Here, we present conserved and liverwort-specific miRNAs and their putative targets that are differentially expressed in vegetative and reproductive Marchantia organs. For four conserved miRNAs (MpmiR529c, MpmiR319a/b, MpmiR160 and MpmiR166a) it was shown that they are essential in reproduction (Tsuzuki et al., 2016; Flores-Sandoval et al., 2018a; Flores-Sandoval et al., 2018b; Tsuzuki et al., 2019). Differential expression pattern of other conserved- and selected liverwort-specific miRNAs may suggest that they also play an important role in various developmental processes, including sexual reproduction. Moreover, we show that all these miRNAs exhibit reverse correlation with the expression of their cognate targets.

## Material and methods

### Plant material and growth conditions

The NGS sequencing and northern blot hybridization experiments were performed using *Marchantia polymorpha L.* plants collected in Chalin (near Sierakow, Poland). *M. polymorpha L.* male accession Takaragaike-1 (Tak-1) and female accession Takaragaike-2 (Tak-2) were used for performing 5’RLM RACE, 3’ RACE, and RT-qPCR experiments. Plants were subjected to a light-to-dark cycle with 16h of light followed by 8h of darkness. The light was provided by LED Neonica Growy (#TSG300; Neonica Poland) at an intensity of 50-60 µmol m^-2^ s^-1^, while maintaining a temperature of 22°C. Plants were grown in axenic cultures on half-strength B5 (Gamborg) medium supplemented with 1% (w/v) sucrose. Additionally, they were also grown on Jiffy-7® (Jiffy International AS) Ø 42mm peat pellets. To induce gametangiophore development, thalli were transferred to high-power blue irradiation (470nm) at a level of 30 µmol m^-2^ s^-1^, deep red irradiation (660nm) at a level of 30 µmol m^-2^ s^-1^, and far-red irradiation (735-740nm_ at a range of 20-30 µmol m^-2^ s^-1^ emitted from diodes sourced from LED Engin (OSRAM GmbH).

### Extraction of total RNA

Total RNA for sRNA NGS sequencing and for northern blot experiments was isolated using a method that enables the enrichment of small RNAs (Kruszka et al., 2013). RNA was extracted from following Marchantia organs: antheridiophores, archegoniophores, male and female vegetative thalli, in three biological repetitions. For RT-qPCR and RACE experiments, Direct-zol^TM^ RNA Mini-prep kit (Zymo Research) was used according to manufacturer’s protocol. In this case, 10 µg of RNA was subjected for DNase I treatment using TURBO DNA-free^TM^ kit (Ambion^TM^; Thermo Fisher Scientific) according to manufacturer’s protocol.

### 5’RLM RACE and 3’RACE

For 5’ RLM and 3’-RACE, 2-5 µg of each DNase I treated total RNA was used for cDNA synthesis and experiments were performed using GeneRacer^TM^ kit (# L150201; Invitrogen™; Thermo Fisher Scientific) according to manufacturer’s instructions, with several modifications (Knop et al., 2017). For *MIR* gene structure analysis, MpTAK v6.1 was used as a reference genome (Bowman et al., 2017; Montgomery et al., 2020).

### RT-qPCR

For the detection and quantification of pri-miR and miRNA target levels, we performed RT-qPCR analysis using Sso Advanced Universal SYBR Green Supermix (BioRad) and the 7900 HT Fast-Real time PCR system (Thermo Fisher Scientific). Three biological replicates were used for each sample. cDNA templates were synthesized by reverse transcription from total RNA using SuperScript^TM^ III reverse transcriptase (200 U/µl), according to manufacturer’s protocol. MpACTIN 7 was chosen as a reference gene (Saint-Marcoux et al., 2015). All primers used in the experiments are listed in Table S1. The statistical significance of the results presented was estimated using a student’s t-test at three significance levels: *p < 0.05, **p < 0.01, and ***p < 0.001.

### Degradome library preparation and data analysis

Construction of Parallel Analysis of RNA Ends (PARE; RNA degradome) libraries was done for the study of cleaved mRNA targets. Libraries were prepared from poly(A)-enriched RNA from following Marchantia organs: vegetative female thalli, archegoniophores, vegetative male thalli and antheridiophores. Preparation of each library was like previously described in the protocols (Addo-Quaye et al., 2009; German et al., 2009) with slight modifications (Grabowska et al., 2020; Sega et al., 2021). All RNA and DNA adapters were purified by PAGE electrophoresis. PCR reactions were performed in a 50 μl volume containing Q5® Hot Start High-Fidelity DNA Polymerase (New England Biolabs). Quantitative analysis of the purified libraries was performed using Qubit 3.0 Fluorometer and Qubit® dsDNA HS Assay kit (Invitrogen^TM^). The four libraries were pooled together in equal molar ratio and sequenced by Fasteris SA company (Switzerland). Bioinformatic analysis of the degradome data followed the methodology outlined in (Alaba et al., 2015). Identification of target mRNAs employed specific parameters, including the degradome score, raw and normalized reads, and the position in the ranked cleavage sites of a given cDNA. The degradome score was based on cutting power and compliance with values ranging from 0 to 18, with negative points assigned for sequence mismatches.

### sRNA sequencing and data analysis

5µg of total RNA enriched in sRNA fraction was isolated from antheridiophores, archegoniophores and male and female vegetative thalli, respectively. The isolation was performed in three biological repetitions, according to the protocol published by (Kruszka et al., 2013). RNA integrity number (RIN) was measured using 2100 Bioanalyzer Instrument and Agilent RNA 6000 Nano Kit (Agilent Technologies). RNA samples with the RIN scale ranges from 9 to 10 were used. NGS sequencing was performed using Illumina HiSeq 2500 instrument, with the number of cycles being 1 × 50 + 7, using HiSeq SBS Kit v4 (Illumina) (Fasteris; Switzerland). Adapter sequences were eliminated utilizing the cutadapt program (version 4.1) following the procedure outlined in (Martin, 2011). Reads ranging from 18 to 28 nucleotides in length were chosen for subsequent analysis. The fastx_collapser function from the FASTX-toolkit (version 0.0.14) package was employed to count the reads. Additional analyses, including RPM normalization, were conducted using the R programming environment.

### Northern blot hybridization analysis of mature miRNA

Northern blot experiments were performed as described in (Kruszka et al., 2013). Up to 15 μg of total RNA per line was loaded on denaturing PAGE gels. A probe complementary to U6 snRNA was used as a loading control (Szarzynska et al., 2009). The oligonucleotide sequences that were used as probes are shown in Table S1. During our studies on miRNA accumulation using sRNA NGS sequencing data and northern blot hybridization experiments in all organs studied generally we obtained identical results. However, in few cases we encountered discrepancies in both techniques. The reason for these discrepancies is not clear for us. In these ambiguous cases, we decided to perform northern blot hybridization experiments multiple times, and since we always obtained the same results, these data were treated as more reliable.

## Results

### Conserved miRNAs are differentially expressed in M. polymorpha vegetative and reproductive organs

*M. polymorpha* shares nine miRNA families with other land plants. Targets for these miRNAs in Marchantia have been already designated (Lin et al., 2016; Tsuzuki et al., 2016; Lin & Bowman, 2018; Pietrykowska et al., 2022). We analyzed the expression of selected and the most abundant conserved miRNA family members in male and female vegetative and reproductive organs: antheridiophores and archegoniophores, using sRNA NGS sequencing and northern blot hybridization experiments. All tested miRNAs exhibit different accumulation patterns in vegetative and reproductive organs (Fig.1 A-F).

**Figure 1.**
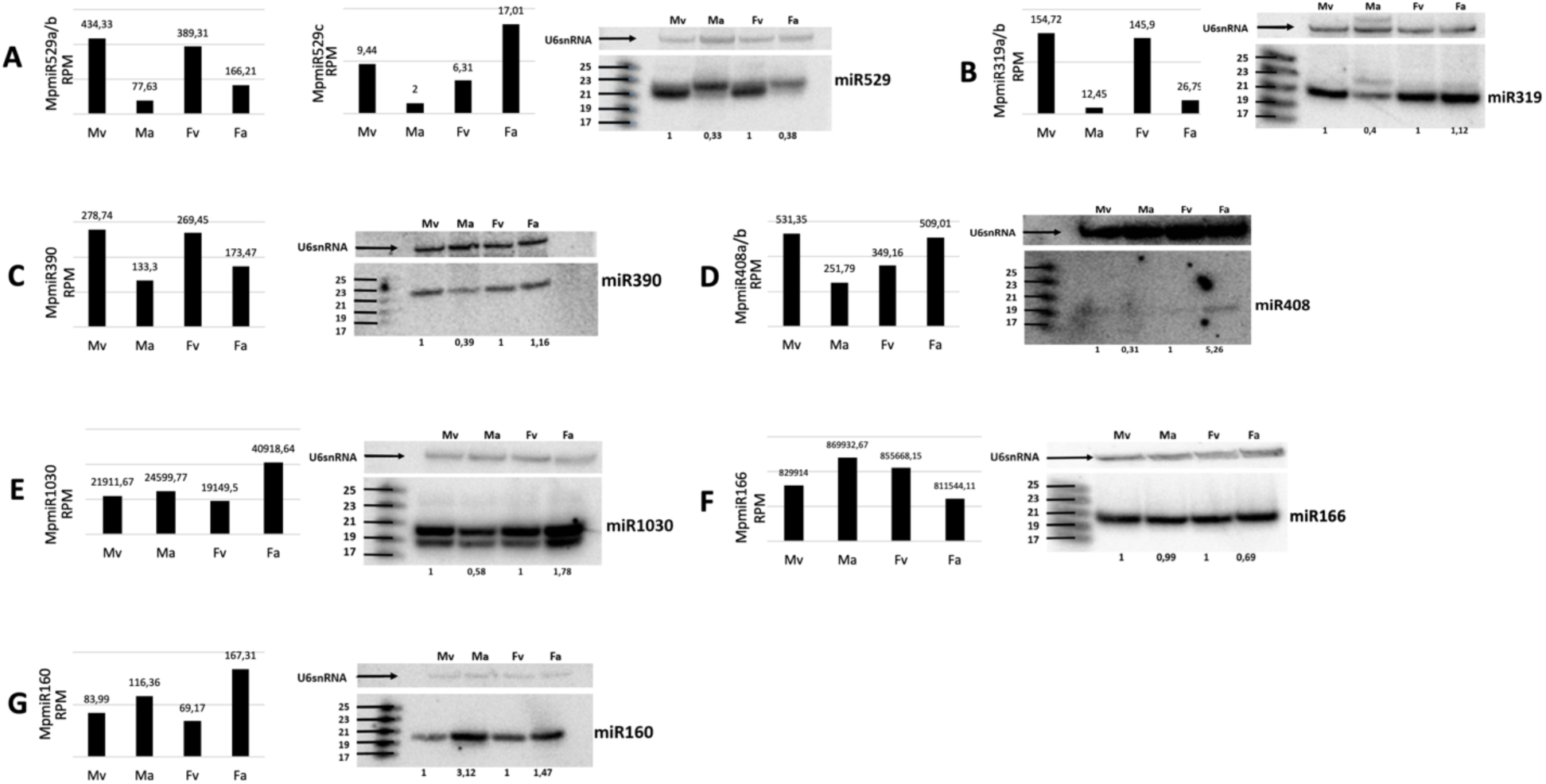
Accumulation of conserved miRNAs in *M. polymorpha* vegetative and reproductive organs. (A-G) sRNA NGS data (left panels) and northern blot hybridization results (right panels) from male vegetative thalli (Mv), antheridiophores (Ma), female vegetative thalli (Fv) and archegoniophores (Fa); normalized read counts are presented above each bar; RPM-reads per million; U6snRNA was used as RNA loading control; the numbers below blot images are the relative intensities of miRNA bands; control signals in each blot were treated as 1; differences in signal intensity were calculated separately for male vegetative thalli control/antheridiophores, female vegetative thalli control/archegoniophores; the left side of northern blots shows the RNA marker depicting 17-25 nucleotide long RNAs

We found that all MpmiR529 family members show downregulation in reproductive organs in Marchantia. Specifically, sRNA NGS data revealed that MpmiR529a/b is downregulated in both: antheridiophores and archegoniophores, and MpmiR529c is downregulated in antheridiophores but upregulated in archegoniophores. Overall, the expression level of MpmiR529c is strongly downregulated when compared to MpmiR529a/b (Fig.1A).

The levels of MpmiR319a/b and MpmiR390 are downregulated in antheridiophores as proved by sRNA NGS sequencing and northern blot hybridization (Fig. 1 B, C). However, NGS data show also strong downregulation of both MpmiR319a/b and MpmiR390 in archegoniophores, but northern blot hybridization shows no difference in the accumulation of these miRNAs when female vegetative thalli and archegoniophores are compared.

MpmiR408a/b and MpmiR1030 exhibit strong upregulation in archegoniophores (Fig. 1D, E). MpmiR408a/b shows downregulation in antheridiophores (Fig. 1D), and the same is observed for MpmiR1030 when northern blot hybridization results are concerned. However, sRNA NGS results demonstrated slight upregulation of MpmiR1030 in antheridiophores (Fig. 1E).

According to sRNA NGS results, there is upregulation of MpmiR166 in antheridiophores and downregulation in archegoniophores (Fig.1F). Northern blot hybridization confirms sRNA NGS results in the case of archegoniophores, however in the case of antheridiophores it shows no difference in expression of MpmiR166 when male vegetative and male generative thalli are compared.

Finally, MpmiR160 is upregulated in Marchantia antheridiophores and archegoniophores as compared to vegetative thalli when sRNA NGS data and northern blot hybridization are considered (Fig.1G).

We compared the expression pattern of described miRNAs and their cognate, conserved and non-conserved targets by using available transcriptomic data from male vegetative thalli, antheridiophores and archegoniophores (Montgomery et al., 2020; Tan et al., 2023). Generally, the reverse correlation between miRNA and their target levels was observed (Fig. S1A-G) (Bowman et al., 2017; Lin & Bowman, 2018; Montgomery et al., 2020). For four out of seven selected conserved miRNA family members, the function in Marchantia development and sexual reproduction has been already proven (Tsuzuki et al., 2016; Flores-Sandoval et al., 2018b; Tsuzuki et al., 2019). Our results clearly suggest that different miRNA accumulation pattern in male and female vegetative thalli, antheridiophores and archegoniophores could be indicative for their involvement in Marchantia development and reproductive processes. The observed relationship encouraged us to study also selected liverwort-specific miRNAs for their different accumulation in vegetative and reproductive organs.

### Gene structures of selected liverwort-specific miRNAs and distinct accumulation of mature miRNAs and their targets in vegetative and reproductive organs

For our analysis, we selected miRNA families described as common to *P. endiviifolia* and *M. polymorpha*. We focused on Pen-miR8163/MpmiR11737a/b and Pen-miR8170/MpmiR11865* (Tsuzuki et al., 2016; Bowman et al., 2017; Pietrykowska et al., 2022). Additionally, we decided to characterize selected liverwort-specific miRNAs that are not present in Pellia but exhibit differential expression profiles in Marchantia, namely MpmiR11887 and MpmiR11796.

### MpmiR11737a/b show different accumulation patterns in Marchantia

sRNA NGS data and northern blot hybridization results showed higher accumulation of MpmiR11737a in male and female vegetative thalli when compared to Marchantia reproductive organs (Fig. 2A, B). Genomic database analysis revealed that MpmiR11737a is encoded within the 2^nd^ intron of *Mp5g12920* gene which encodes chloroplast PsbP protein (Tsuzuki et al., 2016; Bowman et al., 2017; Fig. 2E). Pre-miR11737a forms a classical stem-loop structure (Fig. 2D). Using 5’-RLM and 3’-RACE experiments, we extended the sequence of pre-miR11737a and confirmed that the full-length transcript of *MIR11737a* gene is 1131 nt in length. Its transcription start site (TSS) is located within the intron and it terminates in the 3^rd^ exon of its host gene (Fig. 2E). Hence, it can be concluded that *MIR11737a* represents an independent transcriptional unit which overlaps with the *Mp5g12920* gene sequence. RT-qPCR analysis revealed that the expression profile of pri-MpmiR11737a and mature MpmiR11737a follow the same pattern (Fig. 2C).

**Figure 2.**
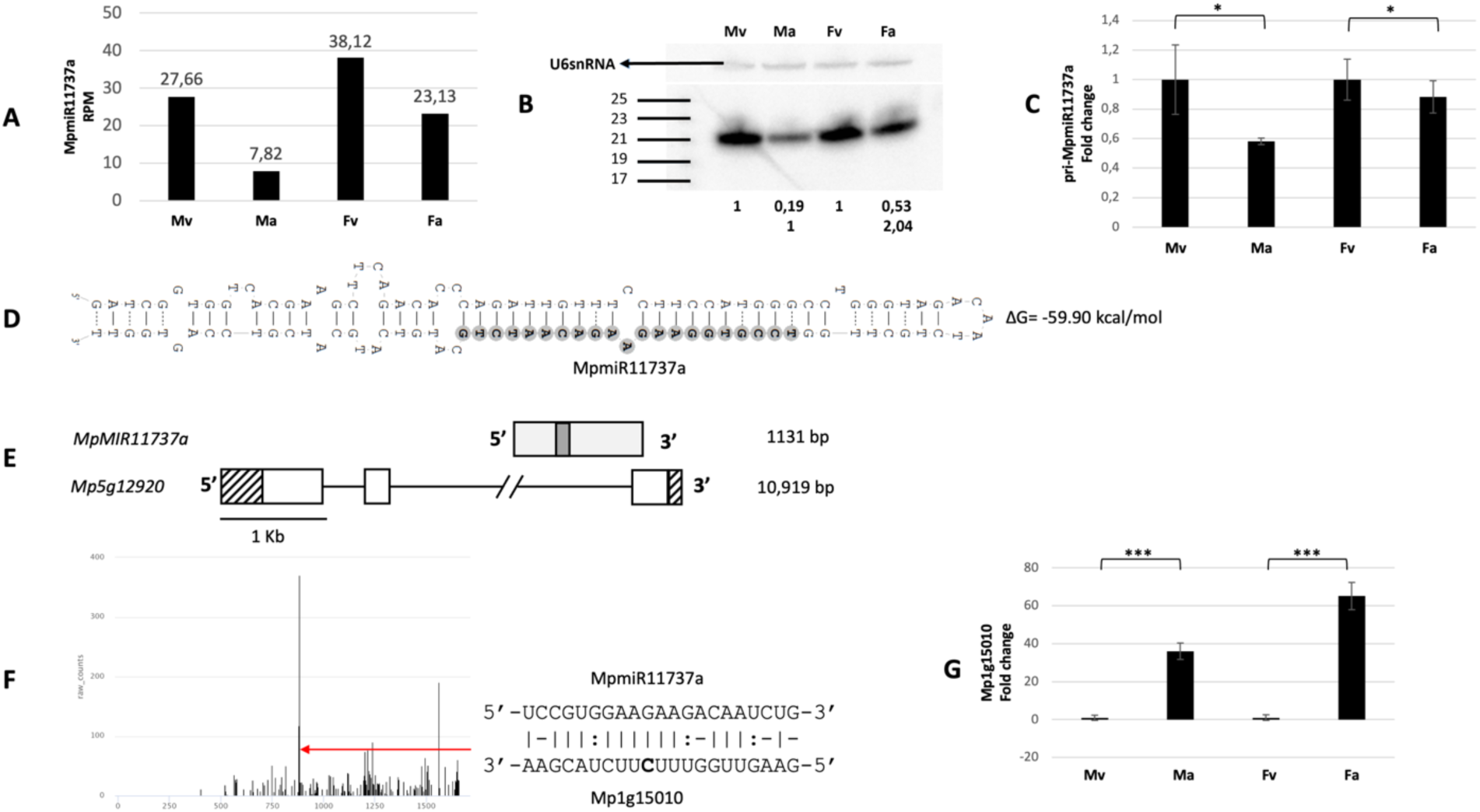
Accumulation profile of liverwort-specific MpmiR11737a, its pri-miRNA level, gene structure and target analyses. (A) sRNA NGS sequencing results; normalized read counts are presented above each bar; RPM-reads per million; (B) Northern blot hybridization analysis; U6snRNA was used as RNA loading control; the numbers below blot images are the relative intensities of miRNA bands; control signals in each blot were treated as 1; differences in signal intensity were calculated separately for male vegetative thalli control/antheridiophores, female vegetative thalli control/archegoniophores (numbers in the first row) or antheridiophores control/archegoniophores (numbers in the second row); the left side of northern blots shows the RNA marker depicting 17-25 nucleotide long RNAs; (C) RT-qPCR expression level of pri-MpmiR11737a; *p-value <0.05; (D) Hairpin structure of pre-MpmiR11737a; nucleotides highlighted in gray represent MpmiR11737a sequence; the minimum free energy (ΔG) of predicted structure is shown on the right side (E) *MIR11737a* gene structure (upper panel; pri-miRNA–light gray; pre-miRNA-dark gray), *Mp5g12920* gene overlapping with *MIR11737a* gene (lower panel); boxes – exons (UTRs – striped; CDS – white); lines –introns; scale bar corresponds to 1kb; (F) Target plot (T-plot) of target mRNA *Mp1g15010* based on degradome data; red arrow points to the mRNA cleavage site; T-plot is accompanied by a duplex of miRNA and its target mRNA; nucleotide marked in bold points to the mRNA cleavage site; (G) RT-qPCR expression level of *Mp1g15010*; ***p-value<0.001; male vegetative thalli (Mv), antheridiophores (Ma), female vegetative thalli (Fv) and archegoniophores (Fa)

The selected mRNA target for MpmiR11737a encodes an uncharacterized protein (*Mp1g15010* gene). The putative miRNA slice site is located within 3’-UTR region (Fig. 2F). The level of mRNA target revealed reverse correlation to MpmiR11737a level: its expression is strongly upregulated in male and female reproductive organs (Fig. 2G). Study performed by Montgomery et al. (2020) revealed a second MpmiR11737a target gene, Mp*5g16770,* which encodes a protein belonging to F-box-like domain superfamily. According to available data, mRNA of this gene is also upregulated in reproductive organs (Montgomery et al., 2020).

The level of mature MpmiR11737b (which differs by 2nt substitutions when compared to miR11737a) revealed by sRNA NGS data is extremely low (Fig. 3A) (Pietrykowska et al., 2022). RT-qPCR analysis of pri-miR11737b showed higher expression level of transcript in reproductive organs when compared to vegetative thalli (Fig. 3B). As in the case of pre-MpmiR11737a, it forms a stable stem-loop structure (Fig. 3C). MpmiR11737b is located within 5’-UTR of *Mp8g07030* gene which encodes a protein of unknown function (Fig. 3D) (Bowman et al., 2017). Attempts to perform 5’RLM and 3’RACE were unsuccessful, probably due to very low expression level of the transcript derived from the host gene.

**Figure 3.**
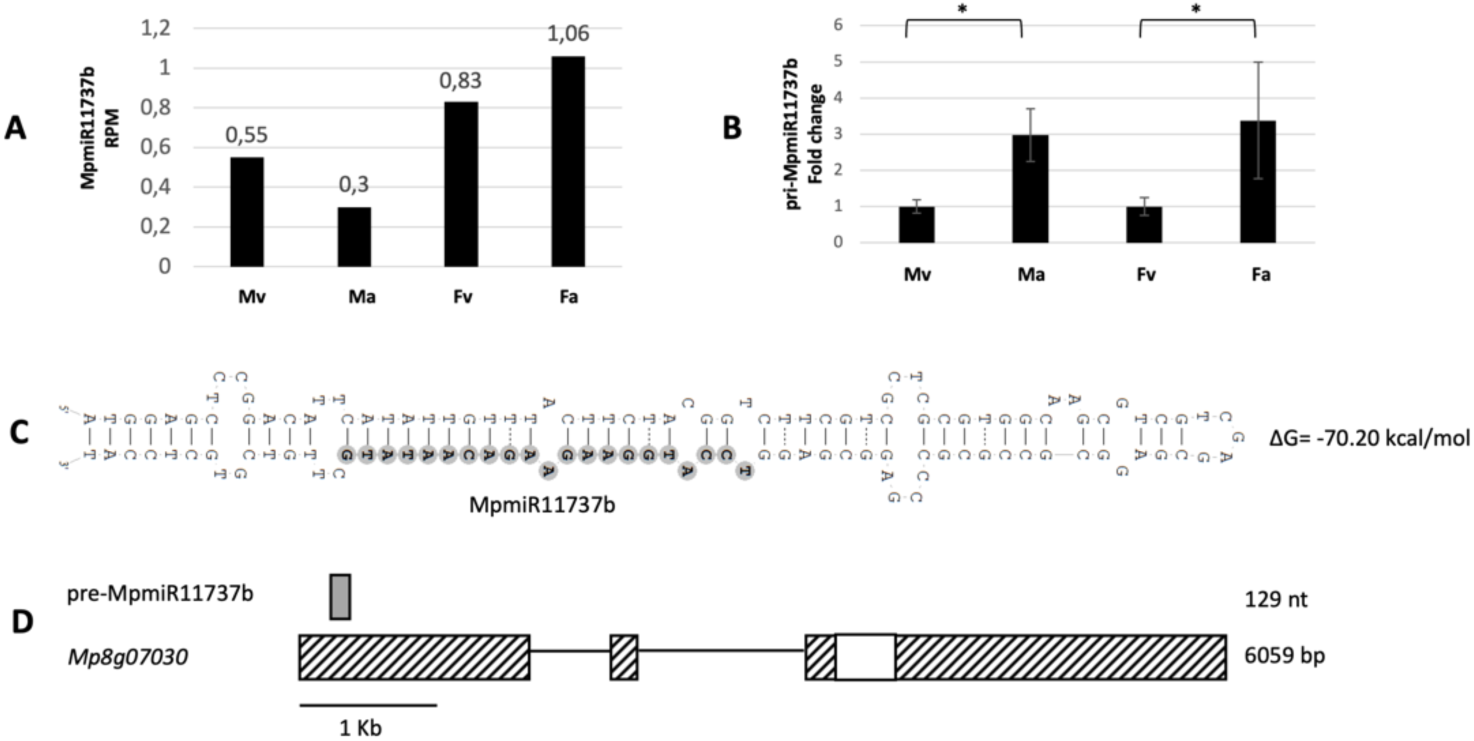
Accumulation profile of liverwort-specific MpmiR11737b, its pri-miRNA level and host gene structure. (A) sRNA NGS sequencing results; normalized read counts are presented above each bar; RPM-reads per million; (B) RT-qPCR expression level of pri-MpmiR11737b; *p-value <0.05; (C) Hairpin structure of pre-MpmiR11737b; nucleotides highlighted in gray represent MpmiR11737b sequence; the minimum free energy (ΔG) of predicted structure is shown on the right side (D) pre-MpmiR11737b (upper panel; dark gray), *Mp8g07030* gene hosting pre-MpmiR11737b (lower panel); boxes – exons (UTRs – striped; CDS – white); lines –introns; scale bar corresponds to 1kb; male vegetative thalli (Mv), antheridiophores (Ma), female vegetative thalli (Fv) and archegoniophores (Fa)

### MpmiR11865 and MpmiR11865* represent two functional miRNA species, originating from the same pri-miRNA

As described earlier, liverwort-specific miRNA, Pen-miR8170, was identified in *M. polymorpha* sRNA NGS data. Surprisingly, it is encoded within the known precursor of MpmiR11865 and represents its passenger miRNA (miRNA*) (Fig. 4A; Tsuzuki et al., 2016; Pietrykowska et al., 2022). Interestingly, miRNA species corresponding to MpmiR11865 was not found in *P. endiviifolia* sRNA NGS data (Alaba et al., 2015). sRNA NGS data showed upregulation of MpmiR11865* in female vegetative thalli and archegoniophores (Fig. 4D). Northern blot hybridization additionally revealed that there is ∼3-fold higher accumulation of MpmiR11865* in archegoniophores than in female vegetative thalli. Unexpectedly, northern blot hybridization revealed that MpmiR11865* accumulates mainly in antheridiophores (Fig. 4D).

**Figure 4.**
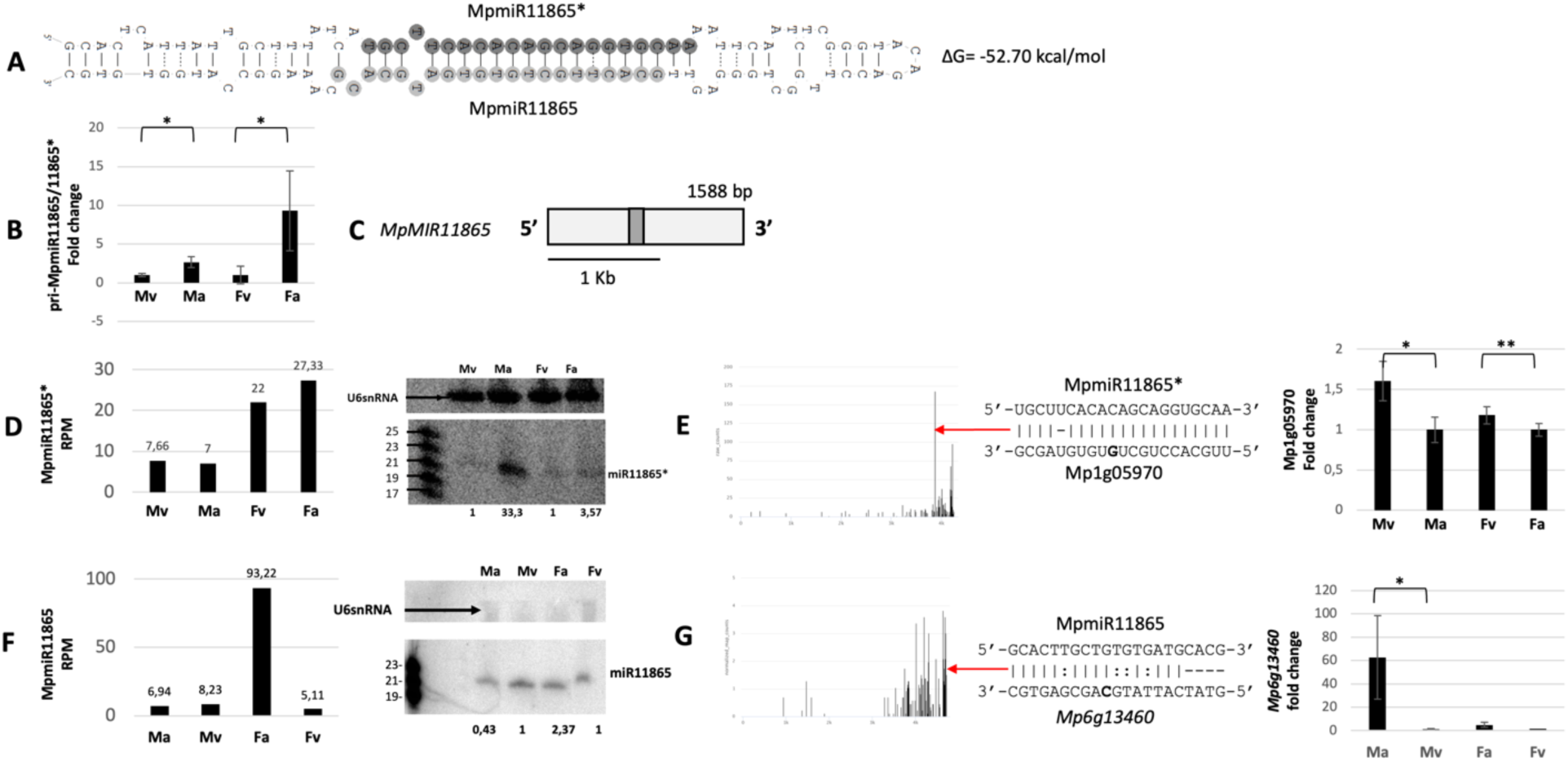
Accumulation profile of liverwort-specific MpmiR11865*/ MpmiR11865, its pri-miRNA level, gene structure and targets analyses. (A) Hairpin structure of pre-MpmiR11865*/ MpmiR11865; nucleotides highlighted in dark gray represent MpmiR11865* sequence and in light gray MpmiR11865 sequence; the minimum free energy (ΔG) of predicted structure is shown on the right side of the structure; (B) RT-qPCR expression level of pri MpmiR11865*/MpmiR11865; *p-value <0.05; (C) Mp*MIR11865* gene structure (pri-miRNA–light gray; pre-miRNA-dark gray); scale bar corresponds to 1kb; (D) MpmiR11865* sRNA NGS sequencing results (left side); normalized read counts are presented above each bar; RPM-reads per million; northern blot hybridization analysis (right side): U6snRNA was used as RNA loading control; the numbers below blot images are the relative intensities of miRNA bands; control signals in each blot were treated as 1; differences in signal intensity were calculated separately for male vegetative thalli control/antheridiophores, female vegetative thalli control/archegoniophores (numbers in the first row) or antheridiophores control/archegoniophores (numbers in the second row); the left side of northern blots shows the RNA marker depicting 17-25 nucleotide long RNAs; (E) Target plot (T-plot) of MpmiR11865* target mRNA (*Mp1g05970)* based on degradome data (left side); red arrow points to the mRNA cleavage site; T-plot is accompanied by a duplex of miRNA and its target mRNA; nucleotide marked in bold points to the mRNA cleavage site; RT-qPCR expression level of *Mp1g05970* (right side); *p-value<0.05; **p-value<0.01; (F) Graph represents MpmiR11865 sRNA NGS sequencing results (left side); normalized read counts are presented above each bar; RPM-reads per million; northern blot hybridization analysis (right side): U6snRNA was used as RNA loading control; the numbers below blot images are the relative intensities of miRNA bands; control signals in each blot were treated as 1; differences in signal intensity were calculated separately for male vegetative thalli control/antheridiophores, female vegetative thalli control/archegoniophores; the left side of northern blot shows the RNA marker depicting 19-23 nucleotide long RNAs; (G) Target plot (T-plot) of MpmiR11865 target mRNA (*Mp6g13460)* based on degradome data (left side); red arrow points to the mRNA cleavage site; T-plot is accompanied by a duplex of miRNA and its target mRNA; nucleotide marked in bold points to the mRNA cleavage site; RT-qPCR expression level of *Mp6g13460* (right side); *p-value<0.05; antheridiophores (Ma), male vegetative thalli (Mv), archegoniophores (Fa) and female vegetative thalli (Fv);

Our studies revealed that mature MpmiR11865 exhibits strong upregulation in archegoniophores (Fig. 4F). Taking into consideration northern results one can conclude that mature MpmiR11865 exhibits opposite expression profile to MpmiR11865*. The level of pri-MpmiR11865 transcript exhibits higher abundance in antheridiophores and archegoniophores, what correlates with northern results showing strong accumulation of MpmiRNA11865* in antheridiophores, and strong accumulation of MpmiR11865 in archegoniophores (Fig. 4 B). Using 5’-RLM and 3’-RACE we found that the full-length transcript of *MIR11865* gene is approximately 1588 nt in length, aligns with the genomic sequence on chromosome 5, and represents an independent intron-less transcriptional unit (Fig. 4C). mRNA of *Mp1g05970* gene, which encodes tRNA ligase1 protein was identified as a potential target of MpmiR11865*. The putative slice site is located within 3’-UTR region and exhibits almost perfect complementarity to MpmiR11865* (Fig. 4E). Moreover, RT-qPCR of identified target mRNA showed reverse correlation to MpmiR11865* level (Fig. 4E). As for MpmiR11865, we identified a potential new target encoding ATPase valosin-containing protein (*Mp6g13460)*. The putative slice site is located within 3’-UTR region and exhibits strong complementarity to MpmiR11865 (Fig. 4G). Its mRNA level also shows reverse correlation to miRNA accumulation (Fig. 4G).

Our studies indicate that both MpmiR11865 and MpmiR11865* represent active miRNA molecules, that undergo different and selective maturation during pri-miRNA11865 processing in male and female reproductive organs. This suggests that both MpmiR11865 and MpmiR11865* can be involved in regulatory mechanisms related to antheridiophores and archegoniophores development and function.

### MpmiR11887 accumulates exclusively in antheridiophores

Our sRNA NGS sequencing data and northern blot hybridization results showed that MpmiR11887 is exclusively expressed in antheridiophores (Fig. 5A, B). RT-qPCR analysis of pri-MpmiR11887 revealed that it is also exclusively detected in antheridiophores (Fig. 5D). Since both, pri-MpmiR11887 and MpmiR11887 are exclusively present in antheridiophores, it suggests transcriptional regulation of MpmiR11887 in male reproductive organs. The exclusive presence of mature MpmiR11887 in antheridiophores suggests its important role in male reproductive organs development.

**Figure 5.**
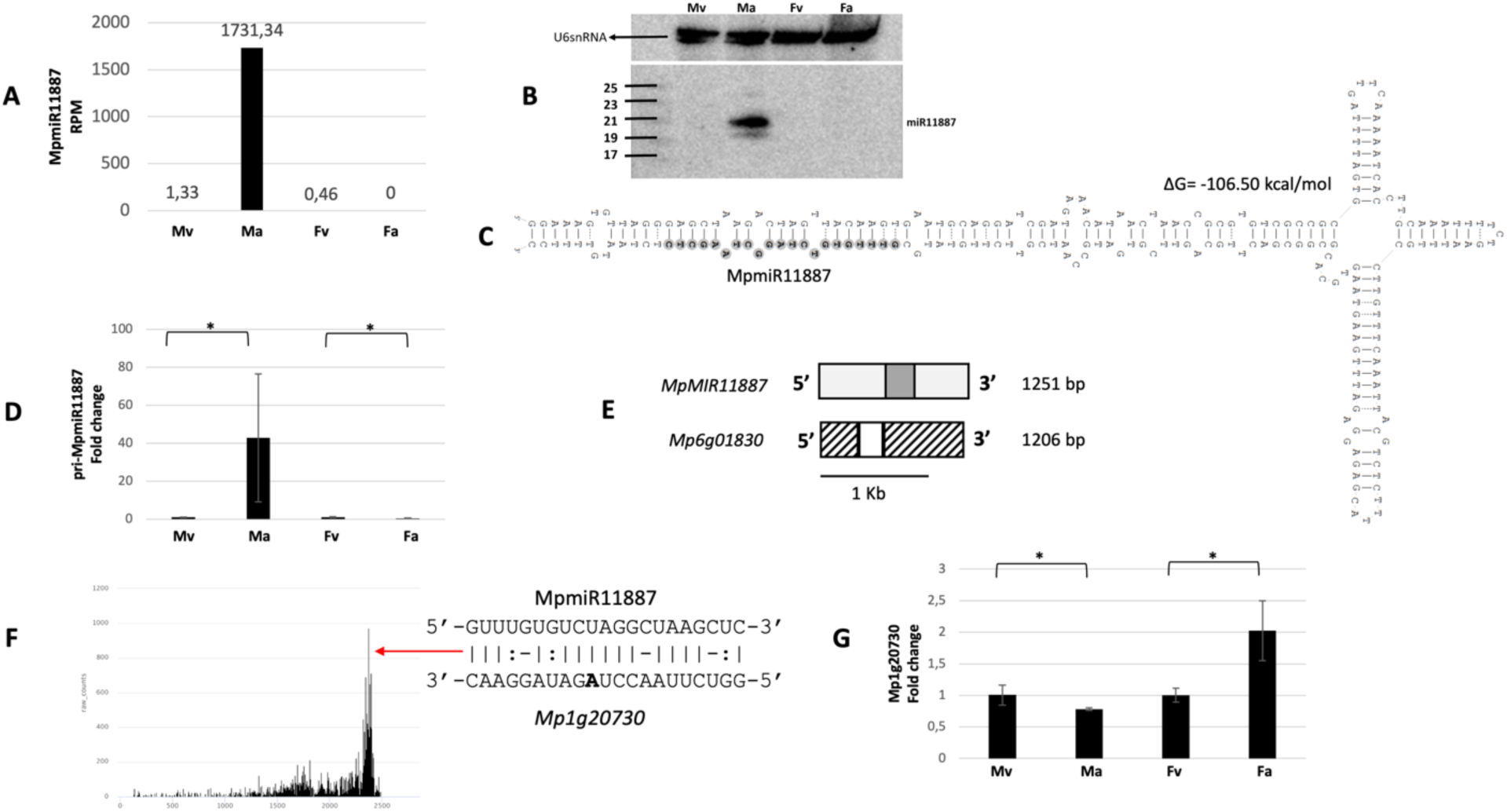
Accumulation profile of liverwort-specific MpmiR11887, its pri-miRNA level, gene structure and target analyses. (A) sRNA NGS sequencing results; normalized read counts are presented above each bar; RPM-reads per million; (B) Northern blot hybridization analysis; U6snRNA was used as RNA loading control; the left side of northern blots shows the RNA marker depicting 17-25 nucleotide long RNAs; (C) Hairpin structure of pre-MpmiR11887; nucleotides highlighted in gray represent MpmiR11887 sequence; the minimum free energy (ΔG) of predicted structure is shown above the precursor; (D) RT-qPCR expression level of pri-MpmiR11887; *p-value <0.05; (E) *MIR11887* gene structure (upper panel; pri-miRNA–light gray; pre-miRNA-dark gray), putative *Mp6g01830* protein-coding gene overlapping with *MIR11887* gene (lower panel); boxes – exons (UTRs – striped; CDS – white); scale bar corresponds to 1kb; (F) Target plot (T-plot) of target mRNA *Mp1g20730* based on degradome data; red arrow points to the mRNA cleavage site; T-plot is accompanied by a duplex of miRNA and its target mRNA; nucleotide marked in bold points to the mRNA cleavage site; (G) RT-qPCR expression level of *Mp1g20730*; *p-value<0.05; male vegetative thalli (Mv), antheridiophores (Ma), female vegetative thalli (Fv) and archegoniophores (Fa)

Analysis of Marchantia genomic database revealed that MpmiR11887 sequence overlaps with 3’-UTR of *Mp6g01830* gene that encodes 65AA long putative protein of unknown function (Bowman et al., 2017). 5’-RLM and 3’-RACE experiments confirmed that the full-length transcript of *MIR11887* gene is 1211 nt long, has the same TSS as the host gene and is 45nt longer (Fig. 5C, E). Degradome sequencing data indicated that MpmiR11887 targets *Mp1g20730* gene mRNA, encoding β-tubulin protein. The putative miRNA slice site is located within 3’-UTR and there is high complementarity between MpmiR11887 and its potential target mRNA (Fig. 5F). The level of putative mRNA target is downregulated in antheridiophores and is reversely correlated to cognate MpmiR11887 level (Fig. 5G).

### MpmiR11796 accumulates mainly in archegoniophores

In the case of another miRNA, MpmiR11796, sRNA NGS and northern blot hybridization results showed its high accumulation in archegoniophores (Fig. 6A, B). RT-qPCR analysis showed lower level of pri-MpmiR11796 in archegoniophores when compared to female vegetative thalli (Fig. 6D). Since mature MpmiR11796 is accumulated mainly in archegoniophores, this suggests post-transcriptional regulation of mature MpmiR11796 level in female reproductive organs. Marchantia genomic database revealed that MpmiR11796 sequence is located within 5’-UTR of *Mp4g11670* gene which encodes a protein of unknown function (Bowman et al., 2017). 5’-RLM and 3’-RACE experiments revealed that *MIR11796* gene represents an independent transcriptional unit which is approximately 505 bp in length and shares sequence overlap with the host gene. The TSS of *MIR11796* gene lies within 5’-UTR of the host gene and 3’-end ends within the first intron (Fig. 6C, E). We identified *Mp4g20750* gene as a potential target for MpmiR11796 (Fig. 6F). It encodes a protein belonging to linker histone H1 and H5 family. RT-qPCR analysis showed exclusive expression of target mRNA in antheridiophores while MpmiR11796 level is the lowest in antheridiophores in comparison to all other organs (Fig. 6G). Interestingly, NGS and northern blot hybridization data show strong accumulation of this miRNA in archegoniophores where histone H1 is not expressed.

**Figure 6.**
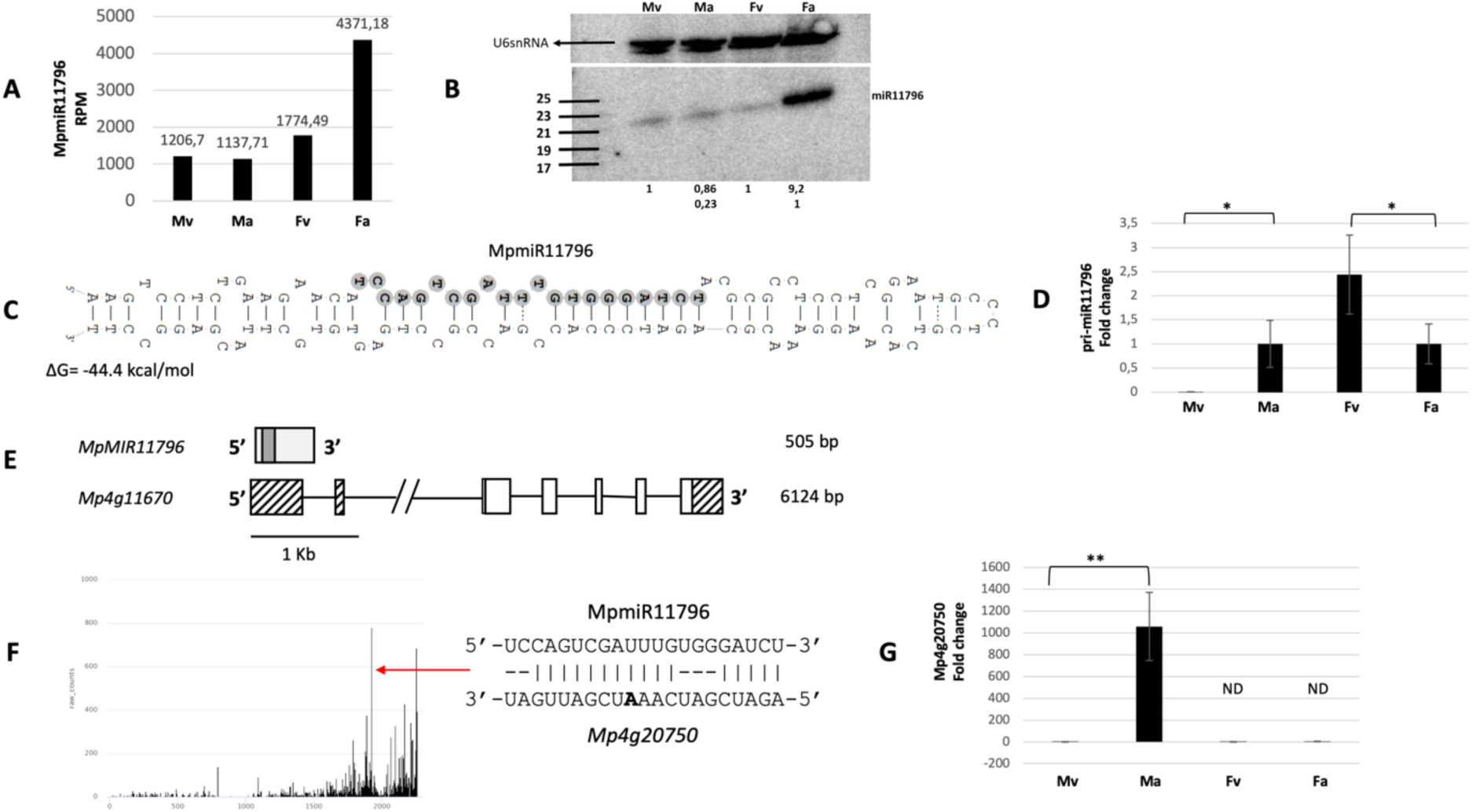
Accumulation profile of liverwort-specific MpmiR11796, its pri-miRNA level, gene structure and target analyses.(A) sRNA NGS sequencing results; normalized read counts are presented above each bar; RPM-reads per million; (B) Northern blot hybridization analysis; U6snRNA was used as RNA loading control; the numbers below blot images are the relative intensities of miRNA bands; control signals in each blot were treated as 1; differences in signal intensity were calculated separately for male vegetative thalli control/antheridiophores, female vegetative thalli control/archegoniophores (numbers in the first row) or antheridiophores control/archegoniophores (numbers in the second row); the left side of northern blots shows the RNA marker depicting 17-25 nucleotide long RNAs; (C) Hairpin structure of pre-MpmiR11796; nucleotides highlighted in gray represent MpmiR11796 sequence; the minimum free energy (ΔG) of predicted structure is shown below the precursor; (D) RT-qPCR expression level of pri-MpmiR11796; *p-value <0.05; (E) *MIR11796* gene structure (upper panel; pri-miRNA–light gray; pre-miRNA-dark gray), *Mp4g11670* gene overlapping with *MIR11796* gene (lower panel); boxes – exons (UTRs – striped; CDS – white); lines –introns; scale bar corresponds to 1kb; (F) Target plot (T-plot) of target mRNA *Mp4g20750* based on degradome data; red arrow points to the mRNA cleavage site; T-plot is accompanied by a duplex of miRNA and its target mRNA; nucleotide marked in bold points to the mRNA cleavage site; (G) RT-qPCR expression level of *Mp4g20750*; **p-value<0.01; male vegetative thalli (Mv), antheridiophores (Ma), female vegetative thalli (Fv) and archegoniophores (Fa)

## Discussion

To date, the biological significance of plant miRNAs has been attributed mostly to conserved miRNAs that control ancestral transcription factors (TFs), or physiological enzymes related to essential plant development and stress tolerance mechanisms (Floyd & Bowman, 2004; Mallory et al., 2005; Palatnik et al., 2007; Sakaguchi & Watanabe, 2012; L. L. Zhang et al., 2022b). Our study revealed that conserved, and selected liverwort-specific miRNAs, as well as their targets are differentially expressed in vegetative and reproductive organs of Marchantia. The unique expression pattern of these miRNAs, along with their inverse correlation to the expression of corresponding targets, indicate their involvement in plant development, the formation of sexual organs and reproductive processes.

Previous characterization of MpmiR529c-*MpSPL2 (SQUAMOSA PROMOTER BINDING PROTEIN-LIKE 2)*, MpmiR319a/b-*MpR2R3-MYB21 (MpMYB33-like* TF), MpmiR166-*MpC3HDZ (HD-ZIPIII TF),* and MpmiR160-*MpARF3 (AUXIN RESPONSE FACTOR 3)* modules showed their important roles in development and sexual reproduction in Marchantia (Floyd & Bowman, 2004; Lin et al., 2016; Tsuzuki et al., 2016; Flores-Sandoval et al., 2018a; Flores-Sandoval et al., 2018b; Tsuzuki et al., 2019). The remaining conserved miRNAs: MpmiR529a/b,

MpmiR408 and MpmiR1030 have not been functionally examined in Marchantia. However, considering their expression profiles observed in vegetative and generative organs, as well as their target levels, it suggests that these modules may be also important for Marchantia development.

There are limited examples of liverwort-specific miRNAs and their targets, whose functional role in the development of Marchantia has been studied. These include FEW RHIZOIDS1 (MpFRH1)- MpRSL1 and Mpo-MR-13–MpSPL1 modules controlling production of rhizoids, gemmae, papillae and meristem dormancy and branching, respectively (Tsuzuki et al., 2016; Honkanen et al., 2018; Thamm et al., 2020; Streubel et al., 2023).

Liverwort-specific MpmiR11865 and MpmiR11865* provide an interesting example of two functional miRNA molecules derived from a common precursor that show opposite accumulation profile in antheridiophores and archegoniophores. This suggests that MpmiR11865 and MpmiR11865* are involved in regulatory mechanisms primarily associated with archegoniophores and antheridiophores development, respectively.

There are reports showing that microRNA* may be also a functional molecule and may accumulate in higher amounts than its microRNA during plant development and stress responses in angiosperms (Meng et al., 2011; Peng et al., 2013; Swida-Barteczka et al., 2023). Thus, *MpMIR11865* gene may represent the first liverwort example of a gene encoding microRNA and microRNA* as functional molecules.

Liverwort-specific MpmiR11887 revealed exclusive expression in antheridiophores. Predicted MpmiR11887 target encodes β-tubulin protein (*Mp1g20730)*. It is known, that in mature pollen of angiosperm specific α- and β-tubulin genes are expressed that are related to pollen tube growth (Carpenter et al., 1992; Kim & An, 1992; Rogers et al., 1993; Villemur et al., 1994; Tchorzewska et al., 2015; Gavazzi et al., 2017). It was also observed, that in liverworts and mosses specific α- and β-tubulin genes were expressed in different tissues (Jost et al., 2004; Sierocka et al., 2011; Buschmann et al., 2016). The expression level of β-tubulin gene (*Mp1g20730)* targeted by MpmiR11887 exhibits lowest expression in antheridiophores. Moreover, data retracted from Marpolbase Expression database show additionally significant downregulation of Mp*1g20730* gene expression in both, antheridia and sperm cells (Montgomery et al., 2020). During Marchantia spermatogenesis a set of processes resulting in remodeling of microtubules and reorganization of endomembrane organelles takes place (Minamino et al., 2022). Thus, MpmiR11887 might be involved in the regulation of spermatogenesis. Interestingly, the full-length transcript of *MIR11887* gene has the same TSS as the proposed protein-encoding host gene. Interestingly, pre-MpmiR11887 is located downstream the short ORF, suggesting that this is *MIRNA* gene encoding miPEP (Fig. 5E). It has been demonstrated that miPEPs are typically encoded by the first open reading frame (miORF) following the TSS located in the 5’ region of the pri-miRNA. MiPEPs have been found to enhance the transcription of pri-miRNAs from which they originate (Lauressergues et al., 2015; Chen et al., 2020; Sharma et al., 2020; Lauressergues et al., 2022).

MpmiR11796 is strongly upregulated in archegoniophores, while its potential target, which encodes a linker histone H1 (*Mp4g20750)* is not detectable. Conversely, this miRNA is weakly accumulating in antheridiophores, while histone H1 mRNA is highly expressed. Based on studies conducted on Arabidopsis, it was demonstrated that there is a significant reduction or even absence in expression of histone H1 during late stages of megaspore mother cell (MMC) development (She et al., 2013; Over & Michaels, 2014). All these data provide a plausible explanation for the reduced level of histone H1 (*Mp4g20750)* transcript in Marchantia archegoniophores simultaneously with the highest accumulation of MpmiR11796.

Arabidopsis linker histones H1 are depleted during microspore development (He et al., 2019). However, in Marchantia, strong upregulation of *Mp4g20750* in antheridiophores and sperm cells is observed (Montgomery et al., 2020; Tan et al., 2023). Further studies are required to confirm the role of MpmiR11796-histone H1 module in Marchantia sexual reproduction.

In angiosperms, miRNAs generally target coding sequence regions (Rhoades et al., 2002; Jones-Rhoades & Bartel, 2004; Ding et al., 2012; Liu et al., 2017). However, in *A. thaliana*, there are several examples of cleavage site localization in the 3’ UTR of mRNA targets: miR156/157- *At1g53160*, miR156-*At2g33810*, miR169-*At1g17590/At1g54160*, and miR319b.2 -*At1g53910* modules (Rhoades et al., 2002; Song et al., 2011; Sobkowiak et al., 2012).

In *M. polymorpha,* like in angiosperms, the miRNA cleavage sites are located mainly in the coding sequence (Lin et al., 2016, Tsuzuki et al., 2016; Honkanen et al., 2018; Thamm et al., 2020; Streubel et al., 2023). However, there are a few examples where the miRNA cleavage sites are in the 3’ UTR. Examples of such cases include MpmiR11667.6-*Mp1g06610* module and MpmiR11670.1-*Mp8g00880* (Lin et al., 2016). In this paper, we propose five additional liverwort-specific miRNA-mRNA modules where the predicted miRNA cleavage sites are within the 3’UTR of the corresponding mRNAs.

Our knowledge on liverwort *MIR* gene structures and pri-miRNA processing is scarce. To understand organ-specific expression of Marchantia miRNAs it is necessary to learn about their gene structures and possible miRNA transcriptional and co/posttranscriptional regulation. Bioinformatic analysis of miRNA-encoding loci in Marchantia genome revealed that majority of the miRNAs are encoded within protein-coding genes (Bowman et al., 2017; Pietrykowska et al., 2022). In this study, we demonstrate that selected *MIR* genes in Marchantia, which were predicted to overlap with protein-coding genes, represent intron-less and independent transcriptional units.

In Arabidopsis, majority of the *MIR* genes also represents independent transcriptional units (Szarzynska et al., 2009; Bielewicz et al., 2012; Zielezinski et al., 2015; Stepien et al., 2017). It would be beneficial to experimentally verify other Marchantia *MIR* gene structures to understand regulatory pathways that affect the final level of mature miRNA.

In angiosperms, it has been observed that the level of miRNAs does not always correlate with the level of pri-miRNA (Barciszewska-Pacak et al., 2015). This observation holds true for Marchantia as well, specifically for pri-MpmiR11865 and pri-MpmiR11796. The lack of correlation may reflect co/post-transcriptional events that affect the final level of mature miRNA. To understand these inconsistencies further studies are required.

Overall, described data on liverwort-specific miRNA-target modules provide a new venue for future research to broaden our knowledge on Marchantia development, sexual reproduction, and control of proper miRNA accumulation during miRNA biogenesis.

## Author contributions

BA, HP performed experiments and participated in manuscript writing, WMK and PN participated in bioinformatic analyses. AJ participated in discussions and writing. ZSK designed experiments and participated in manuscript writing

## Acknowledgements

This research was funded by National Science Centre (NCN): PRELUDIUM 2014/13/N/NZ3/00321, OPUS 2020/39/B/NZ3/00539 and IDUB-Uczelnia Badawcza (05/IDUB/2019/94) at Adam Mickiewicz University, Poznan, Poland.

## Data availability statement

The data that support the findings of this study are available in the National Center for Biotechnology Information (NCBI) Sequence Read Archive database at https://www.ncbi.nlm.nih.gov/bioproject/1008567

---

**Figure S1.**
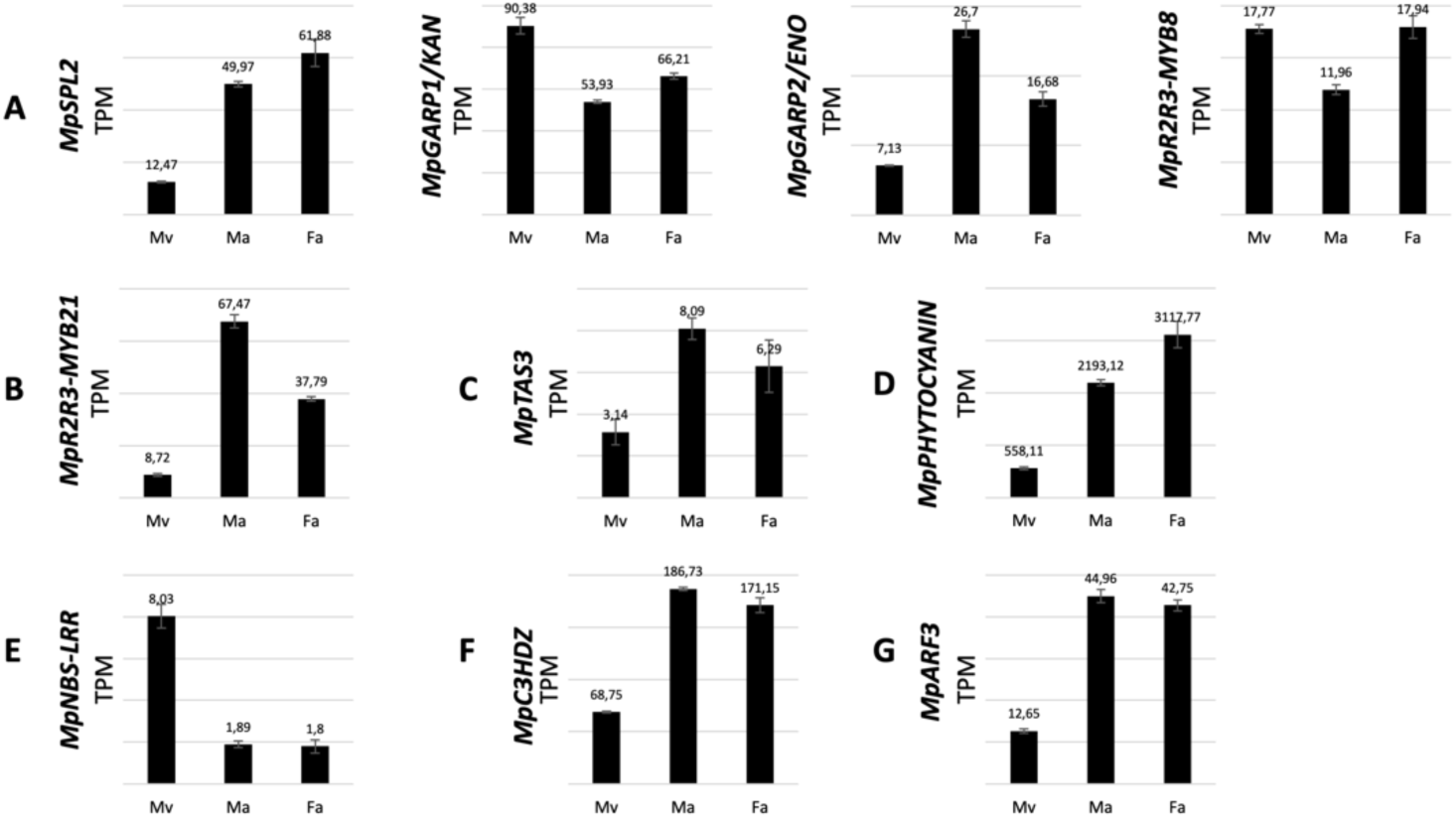
(A-G) Expression profiles of cognate targets for conserved miRNAs in *M. polymorpha* according to Montgomery et al, (2020); (A) targets for MpmiR529c (*MpSPL2)* and MpmiR529a/b (*MpGARP1/KAN*, *MpGARP2/ENO*, *MpR2R3-MYB8*); (B) target for MpmiR319a/b (*MpR2R3-MYB21*); (C) target for MpmiR390 (*MpTAS3*); (D) target for MpmiR408a/b (*MpPHYTOCYANIN*); (E) target for MpmiR1030 (*MpNBS-LRR*); (F) target for MpmiR166 (*MpC3HDZ*); (G) target for MpmiR160 (*MpARF3*); male vegetative thalli (Mv), antheridiophores (Ma), archegoniophores (Fa)

