## Supplementary material for "MiRNAs differentially expressed in vegetative and reproductive organs of *Marchantia polymorpha* – insights into their expression pattern, gene structures and function": Table S1

**Supplementary Table1: List of primers used in the study**

| Name | Sequence (5'→3') | Use |
|---|---|---|
| 5RLM_737a_R1 | CAGGCCCATGGAAGGAAACA | 5'-RLM RACE |
| 5RLM_737a_R2 | GTGACCACCGATCCATAGATG | 5'-RLM RACE |
| 3RACE_737a_R1 | CATCTATGGATCGGTGGTCAC | 3'-RACE |
| 3RACE_737a_R2 | GTAGACAATCTGCTTGGTCCG | 3'-RACE |
| FLT_737a_F1 | GTATACAGTCCTCGCTCTGA | full-length transcript |
| FLT_737a-F2 | CAGTCCTCGCTCTGAAAAGCAC | full-length transcript |
| FLT_737a_R1 | CTGCAAGCGATAATTCTTCCG | full-length transcript |
| FLT_737a_R2 | ATACTATATCAAAAGTGCAGTAGAC | full-length transcript |
| FLT_11865_F1 | ACTGGCCTGCCGCGATCGGA | full-length transcript |
| FLT_11865_F2 | GGAGCGACCTGTTCGATCCTTTGC | full-length transcript |
| 5RLM_11865_R1 | GCAAGTGCAACTCCCAGCCAA | 5'-RLM RACE |
| 5RLM_11865_R2 | GCACCTGCTGTGTGAAGCAT | 5'-RLM RACE |
| 3RACE_11865_R1 | GGATTCTAGTCAGGGCACTCAT | 3'-RACE |
| 3RACE_11865_R2 | CAATCTCGGGTACAGACCTT | 3'-RACE |
| FLT_11865_R1 | CGAGCCATTCACTTTGTAGAT | full-length transcript |
| FLT_11865_R2 | CACGCGTTCGCATTTCTTAACATACG | full-length transcript |
| 3RACE_11887_R1 | AGTAAGTGCACGCGGCGACTT | 3'-RACE |
| 3RACE_11887_R2 | GTATGCGTTTGTGTCTAGGC | 3'-RACE |
| 5RLM_11887_R1 | GTCTTAGCTCCCTAACACTTTCC | 5'-RLM RACE |
| 5RLM_11887_R2 | GCGACACGCGGTTAGTTATG | 5'-RLM RACE |
| FLT_11887_F1 | AGTACCCTTTCGATCGAGGTCA | full-length transcript |
| FLT_11887_R1 | GTGCAGTTCTCCTTCAGTAGGAAGAG | full-length transcript |
| FLT_11887_R2 | CACAGTAACTCGAGGAAGTAC | full-length transcript |
| 5RLM_11796_R1 | AAGGCCTCGTAAGCACACTCA | 5'-RLM RACE |
| 5RLM_11796_R2 | CGAGGAGGCGCGTAGATCCCA | 5'-RLM RACE |
| 3RACE_11796_R1 | GATTTGTGGGATCTACGCGCCT | 3'-RACE |
| 3RACE_11796_R2 | AGATCCCACCGACCGCCTGAGT | 3'-RACE |
| FLT_11796-F | TAAAGAGCAATGCCACTCTCGGG | full-length transcript |
| FLT_11796-R | GCAATTATGCAATGTTCTGTCTGC | full-length transcript |
| pri11737a-F | GATCGGTGGTCACGAAGCTT | RT-qPCR |
| pri11737a-R | AACCACTGGACGATGCATGA | RT-qPCR |
| pri11865-F | GCACTCATTATTGCTTTATC | RT-qPCR |
| pri11865-R | GCACACCATGGCCTTTGCGT | RT-qPCR |
| pri11796-F | AAGTCCTCTGAAGAACATCC | RT-qPCR |
| pri11796-R | AAGGCCTCGTAAGCACACTCA | RT-qPCR |
| pri11887-F | GGAAAGTGTTAGGGAGCTAA | RT-qPCR |
| pri11887-R | GGAAAACATTAGAGAGCTTAGC | RT-qPCR |
| pri11737b-F | ATGGAGCTCCGGACATTCAT | RT-qPCR |
| pri11737b-R | ATGGAGCACGACAAGCATAT | RT-qPCR |
| MpACT-F | AGGCATCTGGTATCCACGAG | RT-qPCR |

| | | |
|---|---|---|
| MpACT-R | ACATGGTCGTTCCTCCAGAC | RT-qPCR |
| Mp1g20730-F | AGCTGACACTACGGTTCTGGT | RT-qPCR |
| Mp1g20730-R | GAGCTTCGACACATTTCACTGCC | RT-qPCR |
| Mp4g20750-F | GGCTGTACTACGTGCGTTTTCG | RT-qPCR |
| Mp4g20750-R | GACTGACGATATCTGTATGAATCG | RT-qPCR |
| Mp1g05970-F | CACTGCCAGCCCTTGTTGATA | RT-qPCR |
| Mp1g05970-R | TGTGGAGGAATGCTGCTTTCA | RT-qPCR |
| Mp1g15010-F | GCTTCGAGTCGTCCTTCATCA | RT-qPCR |
| Mp1g15010-R | GCTCCGGGATCAAAGCCAGCT | RT-qPCR |
| Mp6g13460-F | GTAGTGAGTCTGGCCTAGTGG | RT-qPCR |
| Mp6g13460-R | TCTACACAGTGGTCACAAGGG | RT-qPCR |
| miR160 probe | TGGCATACAGGGAGCCAGGCA | Northern blot |
| miR166 probe | GGGAATGAAGCCTGGTCCGAA | Northern blot |
| miR319ab probe | GGGAGCTCCCTTCAGTCCAAG | Northern blot |
| miR390 probe | GACGCTATCCCTCCTGAGCTT | Northern blot |
| miR408ab probe | AGCCAGGGAAGAGGCAGTGCA | Northern blot |
| miR156a/529 probe | GTGCTCACTCTCTTCTGTCA | Northern blot |
| miR1030 probe | GGTGCAGGTGCAGATGCAGA | Northern blot |
| MpmiR11737ab probe | GAGATTGTTTTCTTCCACGGGG | Northern blot |
| MpmiR11865* probe | ATGCATCTCCTCTGTGAAGCA | Northern blot |
| MpmiR11887 probe | GAGCTTAGCCTAGACACAAAC | Northern blot |
| MpmiR11796 probe | AGATCCCACAAATCGACTGGA | Northern blot |
| U6 probe | TCATCCTTGCGCAGGGGCCA | Northern blot |
